# Integrative and distinctive coding of perceptual and conceptual object features in the ventral visual stream

**DOI:** 10.1101/186924

**Authors:** Chris B Martin, Danielle Douglas, Rachel N Newsome, Louisa LY Man, Morgan D Barense

**Affiliations:** Department of Psychology, University of Toronto, Toronto, ON, Canada; Department of Psychology, Mount Allison University, Sackville, NB, Canada; Rotman Research Institute, Baycrest, Toronto, ON, Canada; Department of Psychology, Queen's University, Kingston, ON, Canada

## Abstract

A tremendous body of research in cognitive neuroscience is aimed at understanding how object concepts are represented in the human brain. However, it remains unknown whether and where the visual and abstract conceptual features that define an object concept are integrated. We addressed this issue by comparing the neural pattern similarities among object-evoked fMRI responses with behavior-based models that independently captured the visual and conceptual similarities among these stimuli. Our results revealed evidence for distinctive coding of visual features in lateral occipital cortex, and conceptual features in the temporal pole and parahippocampal cortex. By contrast, we found evidence for integrative coding of visual and conceptual object features in perirhinal cortex. The neuroanatomical specificity of this effect was highlighted by results from a searchlight analysis. Taken together, our findings suggest that perirhinal cortex uniquely supports the representation of fully-specified object concepts through the integration of their visual and conceptual features.

## Introduction

Semantic memory imbues the world with meaning and shapes our understanding of the relationships among object concepts. Many neurocognitive models of semantic memory incorporate the notion that object concepts are represented in a feature-based manner (Rosch and Mervis, 1975; Tyler and Moss, 2001; Rogers and McLelland, 2004). On this view, our understanding of the concept “hairdryer” is thought to reflect knowledge of observable perceptual properties (e.g., visual form) and abstract conceptual features (e.g., “*used to style hair*”). Importantly, however, there is not always a one-to-one correspondence between how something looks and what it is; a hairdryer and a comb are conceptually similar despite being visually distinct, whereas a hairdryer and a gun are conceptually distinct despite being visually similar. Thus, a fully-specified representation of an object concept requires integration of its perceptual and conceptual features.

Neuroimaging research suggests that object features are stored in the modality-specific cortical regions that supported their processing at the time of acquisition (Thompson-Schill, 2003). However, neurocognitive models of semantic memory differ with respect to how distributed features relate to representations of unified object concepts. On one view, object concepts are thought to emerge through interactions among modality-specific cortical areas (Kiefer and Pulvermüller, 2012; Martin, 2016). Others maintain that they reflect the integration of modality-specific features in trans-modal convergence zones (Damasio, 1989; Rogers et al., 2004; Binder and Desai, 2011), such as the anterior temporal lobes (ATL) (Patterson et al., 2007; Tranel, 2009; Lambon Ralph et al., 2017).

The dominant view of the ATL as a semantic hub was initially shaped by neuropsychological investigations in individuals with semantic dementia (SD) (Patterson et al., 2007). Behaviorally, SD is characterized by the progressive loss of conceptual knowledge across all receptive and expressive modalities (Warrington, 1975; Hodges et al., 1992). At the level of neuropathology, SD is associated with extensive atrophy of the ATL, with the earliest and most pronounced volume loss in the left temporal pole (Mummery et al., 2000; Galton et al., 2001). Most important from a theoretical perspective, patients with SD tend to confuse conceptually similar objects that are visually distinct (e.g., hairdryer – comb), but not visually similar objects that are conceptually distinct (e.g., hairdryer – gun), indicating that the temporal pole expresses conceptual similarity structure (Graham et al., 1994; see Peelen and Caramazza, 2012; Chadwick et al., 2016, for related neuroimaging evidence). Taken together, these findings suggest that the temporal pole supports multi-modal integration of abstract conceptual, but not perceptual, features. Notably, however, a considerable body of research indicates that the temporal pole may not be the only ATL structure that supports feature-based integration.

The representational-hierarchical model of object coding emphasizes a role for perirhinal cortex (PRC), located in the medial ATL, in feature integration that is distinct from that of the temporal pole (Murray and Bussey, 1999). Namely, within this framework PRC is thought to support the integration of conceptual *and* perceptual features. In line with this view, object representations in PRC have been described in terms of conceptual feature conjunctions in studies of semantic memory (Moss et al., 2005; Bruffaerts et al., 2013; Clarke and Tyler, 2014, 2015; Wright et al., 2015), and visual feature conjunctions in studies of visual processing (Barense et al., 2005, 2007; 2012; Lee et al., 2005; Devlin and Price, 2007; Murray et al., 2007; O’Neil et al., 2009; Graham et al., 2010). However, it is difficult to synthesize results from these parallel lines of research, in part, because conceptual and perceptual features tend to vary concomitantly across stimuli (Mur, 2014). For example, demonstrating greater neural pattern similarity in PRC between “horse” and “donkey” than between “horse” and “dolphin” may reflect differences in conceptual or perceptual relatedness. Moreover, because studies linking PRC to the integration of visual features have primarily used pictorial stimuli, it remains unclear whether this result will hold in tasks that require assessment of visual features retrieved from semantic memory. Thus, although the representational-hierarchical account was initially formalized nearly two decades ago (Murray and Bussey, 1999), direct evidence of integration across conceptual and perceptual features remains elusive.

In the current study, we used fMRI to characterize the representational structure of object concepts in the brain. More specifically, we sought to determine whether and where conceptual features are integrated with perceptual features, with an emphasis on visual semantics. This issue was probed, for the first time, using representational similarity analysis (RSA) (Kriegeskorte and Kievit, 2013) and a set of object concepts that were selected to ensure that conceptual similarity was not confounded with visual similarity. In a first step, we generated behavior-based models that captured the conceptual and visual similarities among these object concepts. Next, we scanned participants using task contexts that emphasized processing of either the conceptual or perceptual features of these objects. We hypothesized that both behavior-based models would predict the neural pattern similarities between object concepts, regardless of task context, in brain regions that support the integration of conceptual and perceptual features. Based on the neurocognitive models reviewed above, we anticipated that this result would be uniquely obtained in PRC (Murray and Bussey, 1999; Barense et al., 2011). In addition to PRC, our analysis also probed regions of interest (ROIs) that have been implicated in semantic processing (temporal pole and parahippocampal cortex), and visual processing (lateral occipital cortex) (LOC). We also performed a searchlight analysis, which examined activity patterns within small spheres over the whole brain.

## Results

### Behavior-Based Similarity Models

Using a data-driven approach, we first generated behavior-based models that captured the visual and conceptual similarities among 40 targeted object concepts (Figure 1A-B). Notably, our visual similarity model and conceptual similarity model were derived from behavioral judgments provided by two independent groups of participants. For the purpose of constructing the visual similarity model, the first group of participants (N = 1185) provided pairwise comparative similarity judgments between object concepts (Figure 1A). Specifically, a pair of words was presented on each trial and participants were asked to rate the visual similarity between the object concepts to which they referred using a 5-point Likert scale. Similarity ratings for each pair of object concepts were averaged across participants, normalized, and expressed within a representational dissimilarity matrix (RDM). We refer to this RDM as the *behavior-based visual RDM*.

**Figure 1.**
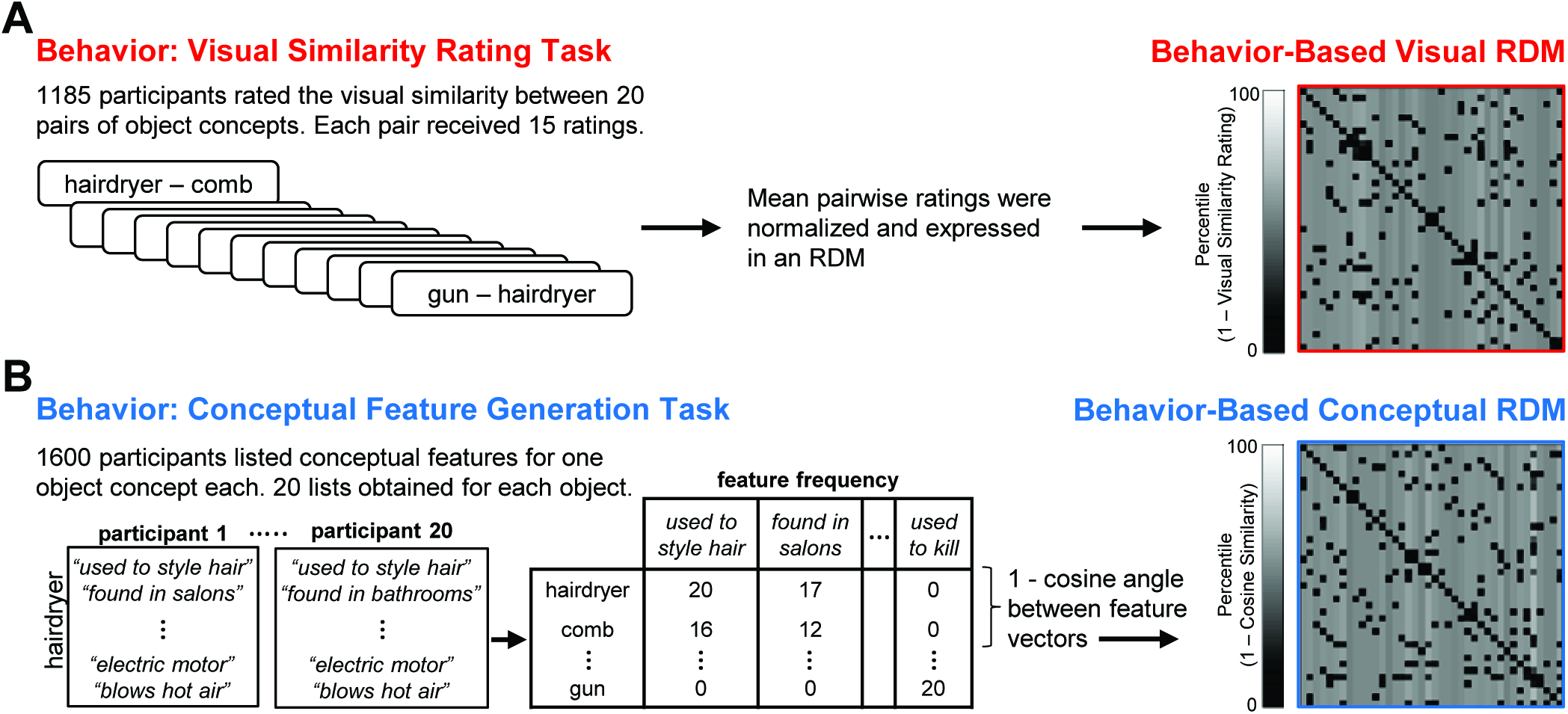
Behavior-based representational dissimilarity matrices (RDMs). **(A)** Visual similarity rating task (left) and corresponding 40x40 behavior-based visual RDM (right). **(B)** Conceptual feature generation task with sample responses from two participants (left), abridged feature matrix depicting the number of participants that listed each feature for each concept (centre), and corresponding 40x40 behavior-based conceptual RDM (right).

For the purpose of constructing the conceptual similarity model, a second group of participants (N = 1600) completed an online feature-generation task (McRae et al., 2005; Taylor et al., 2012) (Figure 1B). Each participant was asked to generate a list of conceptual features that characterize one object concept (e.g., hairdryer: *“used to style hair”, “found in salons”, “electrically powered”, “blows hot air”*; comb: *“used to style hair”, “found in salons”, “has teeth”, “made of plastic”*). Conceptual similarity between all pairs of object concepts was quantified as the cosine angle between the corresponding pairs of feature vectors. With this approach, high cosine similarity between object concepts reflects high conceptual similarity. Cosine similarity values were then expressed within an RDM, which we refer to as the *behavior-based conceptual RDM*.

We next performed a second-level RSA to quantify the relationship between our behavior-based visual RDM and behavior-based conceptual RDM. Critically, this analysis revealed that the model RDMs were not significantly correlated with one another (Kendall’s tau-a = .01, *p* = .09), indicating that differences in visual and conceptual features were not confounded across object concepts. In other words, ensuring that these different types of features varied independently across stimuli (e.g., hairdryer – gun; hairdryer – comb), rather than concomitantly (e.g., horse – donkey; horse – dolphin), allowed us to isolate the separate influence of visual and conceptual features on the representational structure of object concepts in the brain. In this example, a hairdryer and a gun are visually similar but conceptually dissimilar, whereas a hairdryer and a comb are visually dissimilar but conceptually similar.

### fMRI Task and Behavioral Results

We next used fMRI to obtain measurements from which we could infer the representational structure of our 40 object concepts in the neural activity patterns of a third independent group of participants (Figure 2). Functional brain data were acquired over eight experimental runs, each of which consisted of two blocks of stimulus presentation. All 40 object concepts were presented sequentially within each block, for a total of 16 repetitions per concept. On each trial, participants were asked to make a “yes / no” property verification judgment in relation to a block-specific verification probe. Half of the blocks were associated with verification probes that encouraged processing of visual features (e.g., “is the object angular?”), and the other half were associated with verification probes that encouraged processing of conceptual features (e.g., “is the object a tool?”). With this experimental design, we were able to characterize neural responses to object concepts across two task contexts: a visual task context (Figure 2A) and a conceptual task context (Figure 2B).

**Figure 2.**
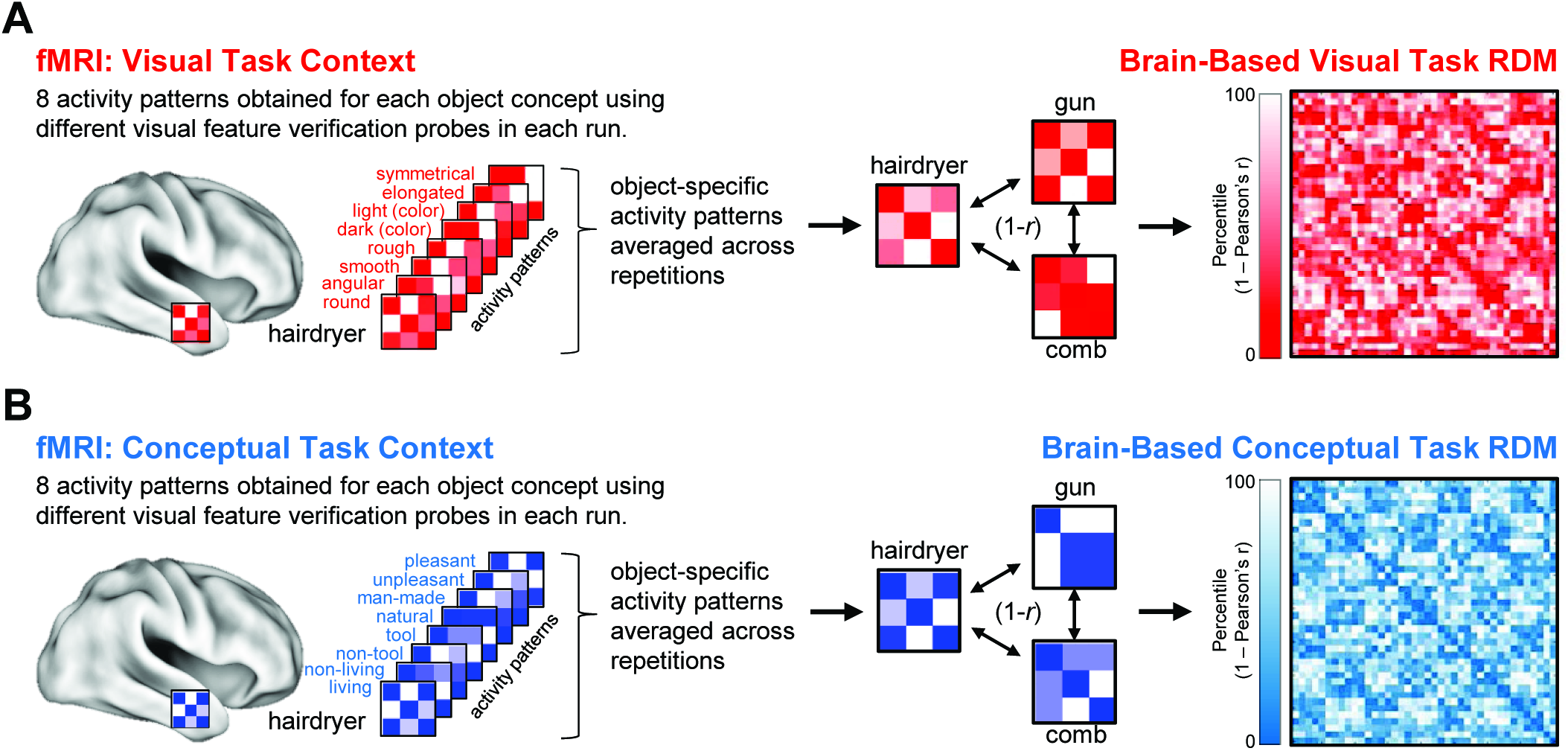
Brain-based representational dissimilarity matrices (RDMs). **(A)** Example of object-evoked neural activity patterns obtained across all eight probes in the visual task context (left), mean object-specific activity patterns averaged across repetitions (center), and corresponding 40x40 brain-based visual task RDM (right). **(B)** Example of object-evoked neural activity patterns obtained across all eight probes in the conceptual task context (left), mean object-specific activity patterns averaged across repetitions (center), and corresponding 40x40 brain-based conceptual task RDM (right).

Behavioral performance on the scanned property verification task indicated that participants interpreted the object concepts and property verification probes with a high degree of consistency (Figure 3). Specifically, all participants (i.e., 16/16) provided the same yes/no response to the property verification task on 88.4% of all trials. Agreement was highest for the “living” verification probe (96.8%) and lowest for the “non-tool” verification probe (73.2%). Moreover, the proportion of trials on which all participants provided the same response did not differ between the visual feature verification task context (Mean = 87.3% collapsed across all eight visual probes) and the conceptual feature verification task context (Mean = 89.5% collapsed across all eight conceptual probes) (*z* = 0.19, *p* = .85). Response latencies were also comparable across the visual feature verification task context (Mean = 1361ms, SD = 302) and the conceptual feature verification task context (Mean = 1388ms, SD = 317) (*t* (15) = 0.61, *p* = .55).

**Figure 3.**
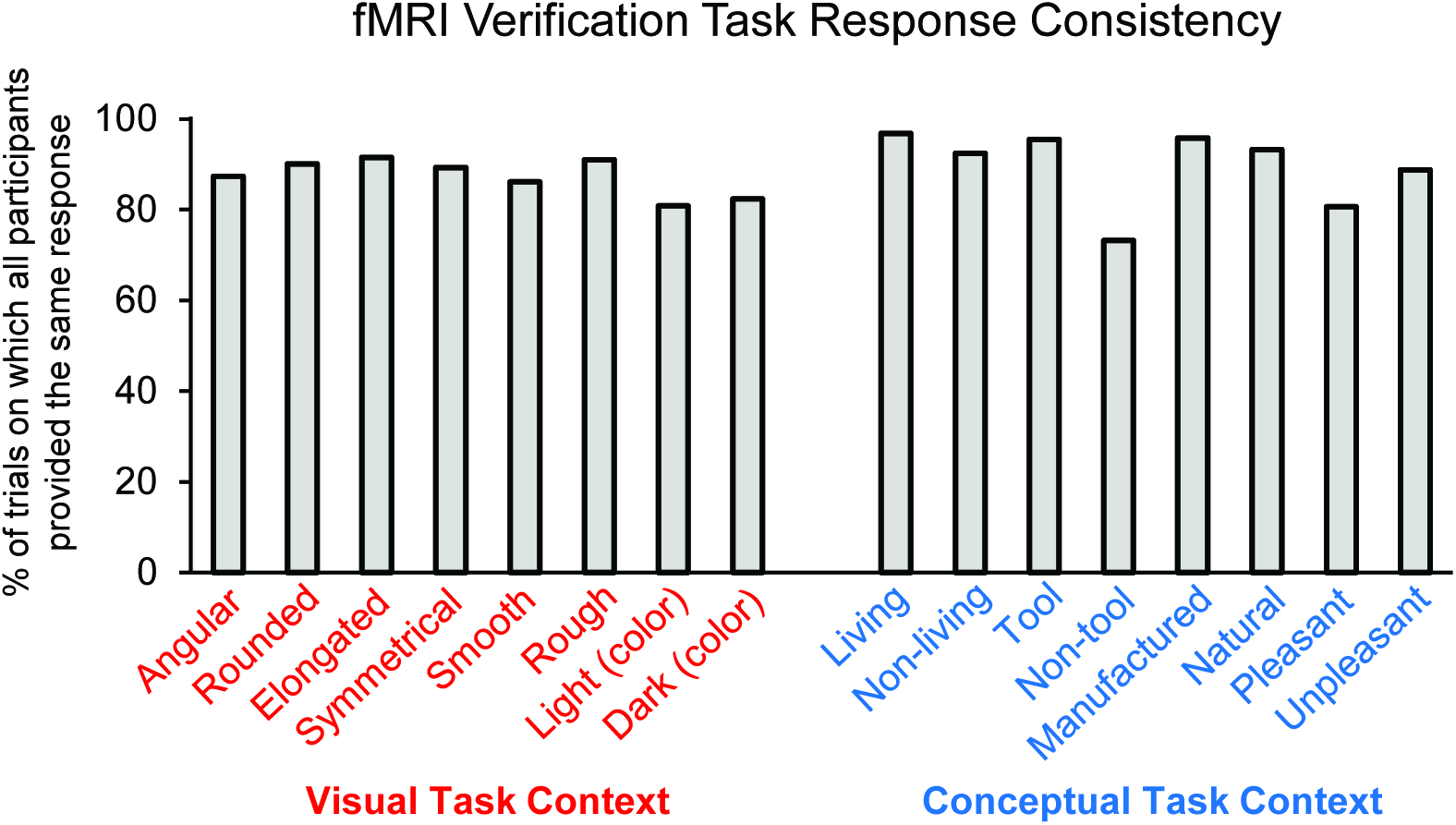
fMRI feature verification task performance. Percentage of trials on which all participants (i.e., 16/16) provided the same ‘yes/no’ response for each property verification probe.

### ROI-Based RSA: Comparison of Behavior-Based RDMs with Brain-Based RDMs

We next quantified pairwise similarities between multi-voxel activity patterns evoked by specific object concepts in the fMRI experiment (Figure 2). For the purpose of conducting ROI-based RSA, we focused on multi-voxel activity patterns obtained in PRC, the temporal pole, parahippocampal cortex, and LOC. ROIs from a representative participant are presented in Figure 4. These ROIs were selected a priori based on empirical evidence linking their respective functional characteristics to visual processing, conceptual processing, or both. Our primary focus was on PRC, which has been linked to integrative coding of visual object features and conceptual object features across parallel lines of research (Barense et al., 2005, 2007, 2012; Lee et al., 2005; O’Neil et al., 2009; Bruffaerts et al., 2013; Clarke and Tyler, 2014; 2015; Wright et al., 2015; Erez et al., 2016). In contrast to PRC, the temporal pole has primarily been linked to processing of conceptual object properties (Mummery et al., 2000; Galton et al., 2001; Patterson et al., 2007; Pobric et al., 2007; Lambon Ralph et al., 2009; Peelen and Caramazza, 2012; Chadwick et al., 2016). A number of studies have also revealed a role for parahippocampal cortex in semantic contextual processing, though its functional contributions remain less well defined than the temporal pole (Bar and Aminoff, 2003, Aminoff et al., 2013, Ranganath and Ritchey, 2012). Lastly, LOC, which is a functionally defined region in occipito-temporal cortex, has been revealed to play a critical role in processing visual form (Grill-Spector et al., 1999; Kourtzi and Kanwisher, 2001; Milner and Goodale, 2006). Because we did not have any a priori predictions regarding hemispheric differences, estimates of neural pattern similarities between object concepts were derived from multi-voxel activity collapsed across the ROIs in the left and right hemisphere.

Mean object-specific multi-voxel activity patterns were estimated in each ROI using general linear models fit to data from the visual and conceptual task contexts, separately. Linear correlation distances (Pearson’s r) were calculated between all pairs of object concepts, which were then expressed in two brain-based RDMs for each ROI. Specifically, the *brain-based visual task RDM* captured the neural pattern similarities obtained between all object concepts in the visual task context (i.e., while participants made visual feature verification judgments) (Figure 2A), and the *brain-based conceptual task RDM* captured the neural pattern similarities obtained between all object concepts in the conceptual task context (i.e., while participants made conceptual feature verification judgments) (Figure 2B).

**Figure 4.**
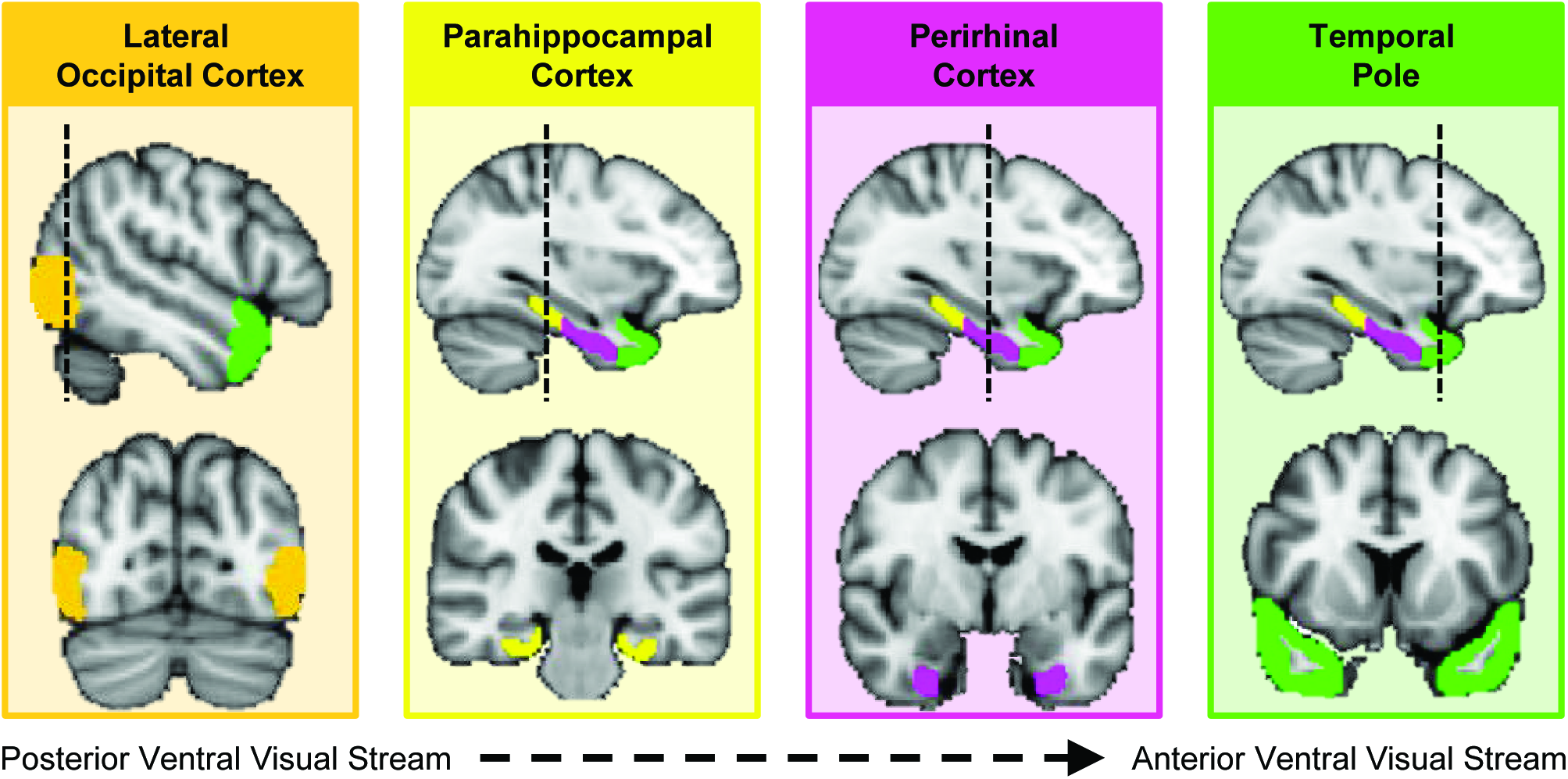
Regions of interest (ROIs) in a representative participant. Cortical regions examined in the ROI-based RSA, including lateral occipital cortex (orange), parahippocampal cortex (yellow), perirhinal cortex (pink), and the temporal pole (green).

We implemented second-level RSA to compare our behavior-based visual and conceptual RDMs (i.e., independent dissimilarity models) with the brain-based visual and conceptual task RDMs (i.e., neural pattern dissimilarity obtained in different verification task contexts) (solid arrows in Figure 5). These analyses were conducted in each ROI using a ranked correlation coefficient (Kendall’s tau-a) as a similarity index (Nili et al., 2014). Significance testing was performed using non-parametric permutation tests for all pertinent comparisons. A Bonferroni correction was applied to compensate for multiple comparisons (4 ROIs x 2 behavior-based RDMs x 2 brain-based RDMs = 16 comparisons, yielding a critical alpha of .003). With this approach, we revealed that object concepts are represented by three distinctive similarity codes that differed across ROIs: visual similarity coding, conceptual similarity coding, and integrative coding. Results from our ROI-based RSA analyses are shown in Figure 6 and discussed in turn below.

**Figure 5.**
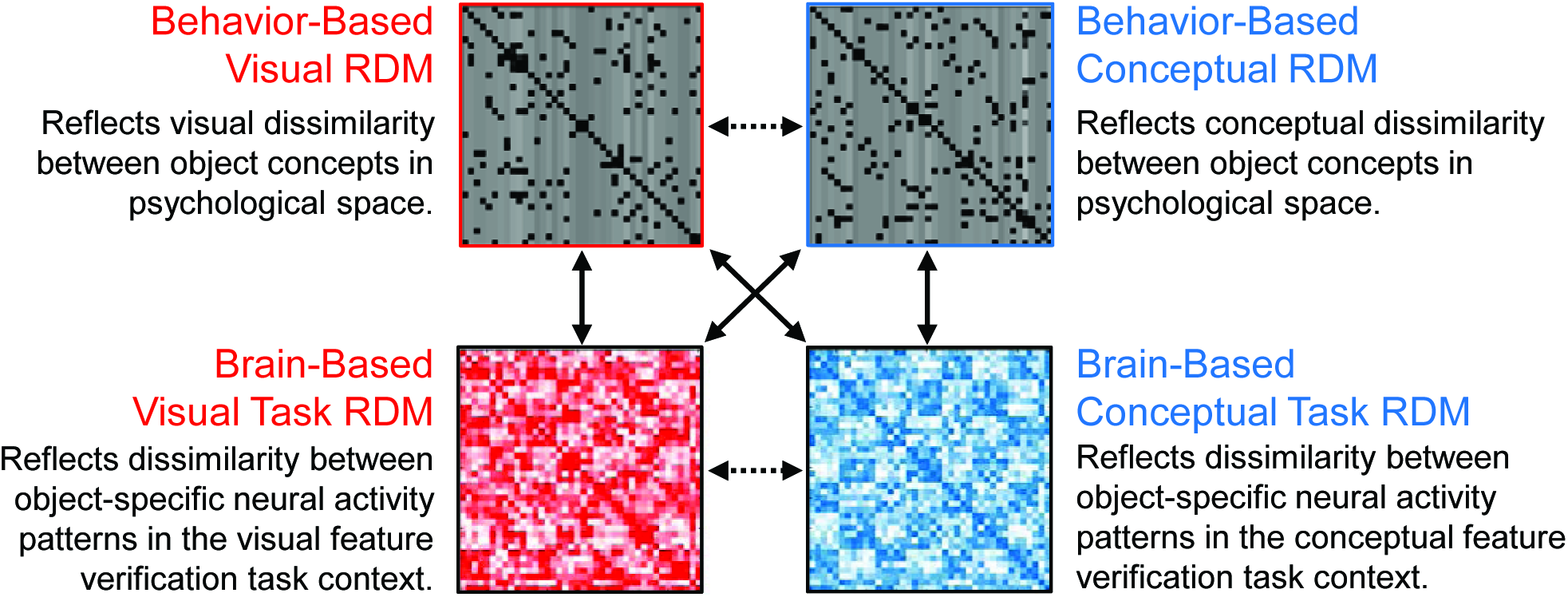
Correlation-based representational similarity analyses (RSA). The dashed horizontal arrow between behavior-based RDMs reflects second-level RSA in which the visual and conceptual models were compared. Solid vertical and diagonal arrows reflect second-level RSA in which behavior-based RDMs were compared with brain-based RDMs. The dashed horizontal arrow between brain-based RDMs reflects second-level RSA in which neural pattern similarities from each task context were directly compared with each other.

**Figure 6.**
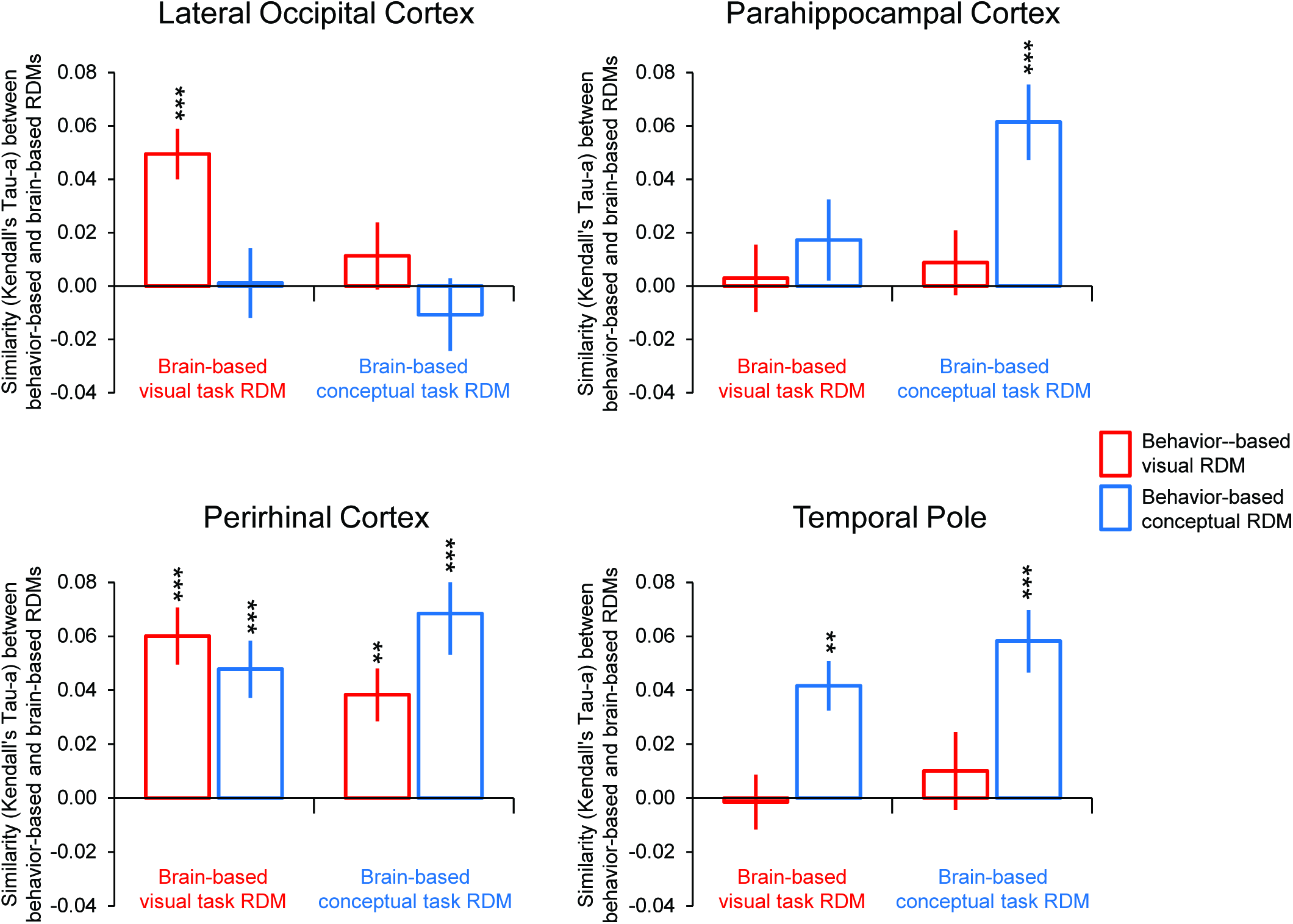
ROI-based RSA results. Similarities between behavior-based and brain-based representational dissimilarity matrices (RDMs) are plotted for each ROI. Similarity was quantified as the ranked correlation coefficient (Kendall’s tau-a) between behavior-based RDMs and the brain-based RDMs. Error bars indicate the standard error, estimated as the standard deviation of 100 deviation estimates obtained from the stimulus-label randomization test. *** *p* < .0001, ** *p* < .001.

#### Lateral Occipital Cortex Represents Object Concepts with a Visual Similarity Code

Consistent with its well-established role in the processing of visual form, patterns of activity within LOC reflected the visual similarity of the object concepts (Figure 6). Specifically, we obtained a significant correlation between the behavior-based visual RDM and the brain-based visual task RDM in LOC (Kendall’s tau-a = .05, *p* < .0001). Notably, however, the correlation between the behavior-based visual RDM and the brain-based conceptual task RDM was not significant (Kendall’s tau-a = .01, *p* = .20). In other words, activity patterns in LOC expressed a visual similarity structure when participants were asked to make explicit judgments about the visual features that characterized object concepts (e.g., whether an object is angular in form), but not when those judgments pertained to features that were conceptual in nature (e.g., whether an object is naturally occurring). Conversely, the behavior-based conceptual RDM did not significantly correlate with the brain-based visual task RDM (Kendall’s tau-a = .002, *p* = .45) or brain-based conceptual task RDM (Kendall’s tau-a = -.016, *p* = .87), indicating that conceptual similarities between object concepts did not capture neural pattern similarities in LOC in either task context. Considered together, these results suggest that LOC represents perceptual information about object concepts in a task-dependent visual similarity code that generalizes across visually similar object concepts that are conceptually distinct (e.g., hairdryer – gun), but not across conceptually similar object concepts that are visually distinct (e.g., hairdryer – comb).

#### The Temporal Pole and Parahippocampal Cortex Represent Object Concepts with a Conceptual Similarity Code

In line with theoretical frameworks that have characterized the temporal pole as a semantic hub (Patterson et al., 2007; Tranel et al., 2009), patterns of activity within this specific ATL structure reflected the conceptual similarity of the object concepts (Figure 6). Specifically, in the temporal pole we revealed a significant correlation between the behavior-based conceptual RDM and the brain-based conceptual task RDM (Kendall’s tau-a = .06, *p* < .0001). The behavior-based conceptual RDM was also significantly correlated with the brain-based visual task RDM (Kendall’s tau-a = .04, *p* < .0001). Thus, the temporal pole expressed a conceptual similarity structure regardless of whether participants were asked to make targeted assessments of conceptual features (e.g., whether the object is a tool) or visual features (e.g., whether it is symmetrical). The behavior-based visual RDM was not significantly correlated with either the brain-based conceptual task RDM (Kendall’s tau-a = .01, *p* = .19) or the brain-based visual task RDM (Kendall’s tau-a = -.001, *p* = .55), suggesting that the representational structure of object concepts in the temporal pole is not shaped by visual properties.

Patterns of activity obtained in parahippocampal cortex, which has previously been associated with the processing of semantically-based contextual associations (Bar and Aminoff, 2003), also reflected the conceptual similarity of the object concepts (Figure 6). Unlike the temporal pole, however, parahippocampal cortex expressed conceptual similarity structure in a task-specific manner. Specifically, the behavior-based conceptual RDM was significantly correlated with the brain-based conceptual task RDM (Kendall’s tau-a = .06, *p* < .0001), but not the brain-based visual task RDM (Kendall’s tau-a = .02, *p* = .10). The behavior-based visual RDM was not a significant predictor of neural dissimilarity structure captured by either the brain-based visual task RDM (Kendall’s tau-a = .002, *p* = .42) or the brain-based conceptual task RDM (Kendall’s tau-a = .009, *p* = .22).

In sum, these results suggest that the temporal pole and parahippocampal cortex represent conceptual information in a manner that enables efficient generalization across conceptually related object concepts that are visually distinct (e.g., hairdryer – comb), but not visually related object concepts that are conceptually distinct (e.g., hairdryer – gun). That is, the degree of similarity between object-evoked activity patterns in these structures reflected the degree of conceptual feature overlap, but not visual feature overlap, between those object concepts. Notably, the temporal pole expressed this conceptual similarity code even when the information that it conveyed was orthogonal to task demands. For example, hairdryer and comb were represented more similarly than were hairdryer and gun, even when task demands encouraged processing of visual features in the visual task context. Conversely, our results suggest that parahippocampal cortex expresses conceptual similarity structure only when task demands prioritize processing of conceptual information in the conceptual feature verification task.

#### Perirhinal Cortex Represents Object Concepts with a Similarity Code that Reflects Integration of Conceptual and Visual Features

Results obtained in PRC support the notion that this structure integrates conceptual and visual object features, as first theorized in the representational-hierarchical model of object representation (Murray and Bussey, 1999). Namely, we revealed that the behavior-based visual RDM and the behavior-based conceptual RDM were each significantly correlated with both the brain-based visual task RDM (behavior-based visual RDM Kendall’s tau-a = .07, *p* < .0001; behavior-based conceptual RDM Kendall’s tau-a = .05, *p* < .0001), and the brain-based conceptual task RDM (behavior-based visual RDM Kendall’s tau-a = .04, *p* < .001; behavior-based conceptual RDM Kendall’s tau-a = .07, *p* < .0001) (Figure 6). These findings indicate that PRC simultaneously expressed both conceptual and visual similarity structure, and did so regardless of whether participants were asked to make targeted assessments of conceptual features (e.g., whether the object concept is living) or visual features (e.g., whether it is elongated). In other words, activity patterns in PRC captured the conceptual similarity between hairdryer and comb, as well as the visual similarity between hairdryer and gun, and did so irrespective of task context. Critically, these results were obtained despite the fact that the brain-based RDMs were orthogonal to one another (i.e., not significantly correlated). Considered together, these results suggest that, of the a priori ROIs considered, PRC represents object concepts at the highest level of specificity through integration of visual and conceptual features.

### ROI-Based RSA: Comparisons of Brain-Based RDMs with Brain-Based RDMs

We next implemented an additional second-level RSA in which we directly compared object-evoked neural similarity patterns within and across our four a priori ROIs. These analyses were conducted using the same methodological procedures applied to compare behavior-based RDMs with brain-based RDMs. We first sought to quantify the representational similarity between the brain-based visual task RDM and brain-based conceptual task RDM obtained within each ROI. This comparison is denoted by the dashed horizontal arrow in the bottom of Figure 5. Notably, these brain-based RDMs were significantly correlated with one another in PRC (Kendall’s tau-a = .06, *p* < .001), but not in the temporal pole (Kendall’s tau-a = .01, *p* = .28), parahippocampal cortex (Kendall’s tau-a = -.01, *p* = .69), or LOC (Kendall’s tau-a = .02, *p* = .18). This result suggests that PRC emphasized similar representational distinctions between object concepts regardless of whether those concepts were processed in the context of a visual or conceptual task context.

In a second set of analyses, we examined whether activity in different ROIs reflected similar representational distinctions across object concepts within the same task context. To this end, we first compared the brain-based visual task RDM obtained in a given ROI with those obtained in all other ROIs. For example, we asked whether the brain-based visual task RDMs obtained in PRC and LOC were significantly correlated with one another for the visual task context. Interestingly, these analyses did not reveal any significant results between any of our ROIs (all Kendall’s tau-a < .029, all *p* > .12). These findings indicate that PRC and LOC, two regions that expressed a visual similarity code as revealed through comparison with the behavior-based visual RDM (Figure 3B), emphasized different visually-based representational distinctions between object concepts.

We next compared the brain-based conceptual task RDM obtained in a given ROI with those obtained in all other ROIs. For example, we asked whether the brain-based conceptual task RDMs obtained in PRC and the temporal pole, were significantly correlated with one another. This set of analyses revealed a trend toward a positive correlation between PRC and parahippocampal cortex (Kendall’s tau-a = .05, *p* < .01, corrected critical alpha = .003), but no such relationship between any other ROIs (all Kendall’s tau-a < .034, all *p* > .08). These findings suggest that although the brain-based conceptual task RDMs obtained in PRC, parahippocampal cortex, and the temporal pole were all significantly correlated with the behavior-based conceptual RDM, they may emphasize different conceptually-based representational distinctions between object concepts.

### Searchlight-Based RSA: Comparisons of Behavior-Based RDMs with Brain-Based RDMs

#### Perirhinal Cortex is the Only Cortical Region that Supports Integrative Coding of Conceptual and Visual Object Features

We next implemented a whole-volume searchlight-based RSA to investigate the neuroanatomical specificity of our ROI-based results. Specifically, we sought to determine whether object representations in PRC expressed visual and conceptual similarity structure within overlapping or distinct populations of voxels. If PRC does indeed support the integrative coding of visual and conceptual object features, then the same subset of voxels in this structure should express both types of similarity codes. If PRC does not support the integrative coding of visual and conceptual object features, then different subsets of voxels should express these different similarity codes. More generally, data-driven searchlight mapping allowed us to explore whether any other regions of the brain showed evidence for integrative coding of visual and conceptual features in a manner comparable to that observed in PRC. To this end we performed searchlight RSA using multi-voxel activity patterns restricted to a 100 voxel ROI that was iteratively swept across the entire cortical surface (Kriegeskorte et al., 2006; Oosterhof et al., 2011). In each searchlight ROI, the behavior-based RDMs were compared with the brain-based RDMs using a procedure identical to that implemented in our ROI-based RSA. These comparisons are depicted by the solid black arrows in Figure 5. The obtained similarity values (Pearson’s r) were mapped to the center of each ROI for each participant separately. With this approach, we obtained participant-specific similarity maps for all comparisons, which were then standardized and subjected to a group-level statistical analysis. A threshold-free cluster enhancement (TFCE) method was used to correct for multiple comparisons with a cluster threshold of *p* < 0.05 (Smith and Nichols, 2009).

All searchlight results are depicted in Figure 7, with corresponding cluster statistics, coordinates, and anatomical labels reported in Table 1. Statistically thresholded group-level similarity maps are presented in Figure 6A for comparison of both behavior-based RDMs with the brain-based visual task RDM, and in Figure 6C for comparison of both behavior-based RDMs with the brain-based conceptual task RDM. To determine whether PRC expressed visual similarity structure and conceptual similarity structure in overlapping or distinct sets of voxels, we examined the extent of voxel overlap across similarity maps. In a first step, we asked whether there were any common voxels across the similarity maps obtained within each task context, separately. Overlapping voxels across similarity maps obtained through comparison of behavior-based RDMs with the brain-based RDM derived from the visual task context are presented in Figure 6B. Overlapping voxels across similarity maps obtained through comparison of behavior-based RDMs with the brain-based RDM derived from the conceptual task context are presented in Figure 6D. Within each task context, we revealed a contiguous cluster of voxels in left PRC in which both behavior-based RDMs predicted task-specific brain-based RDMs.

**Figure 7.**
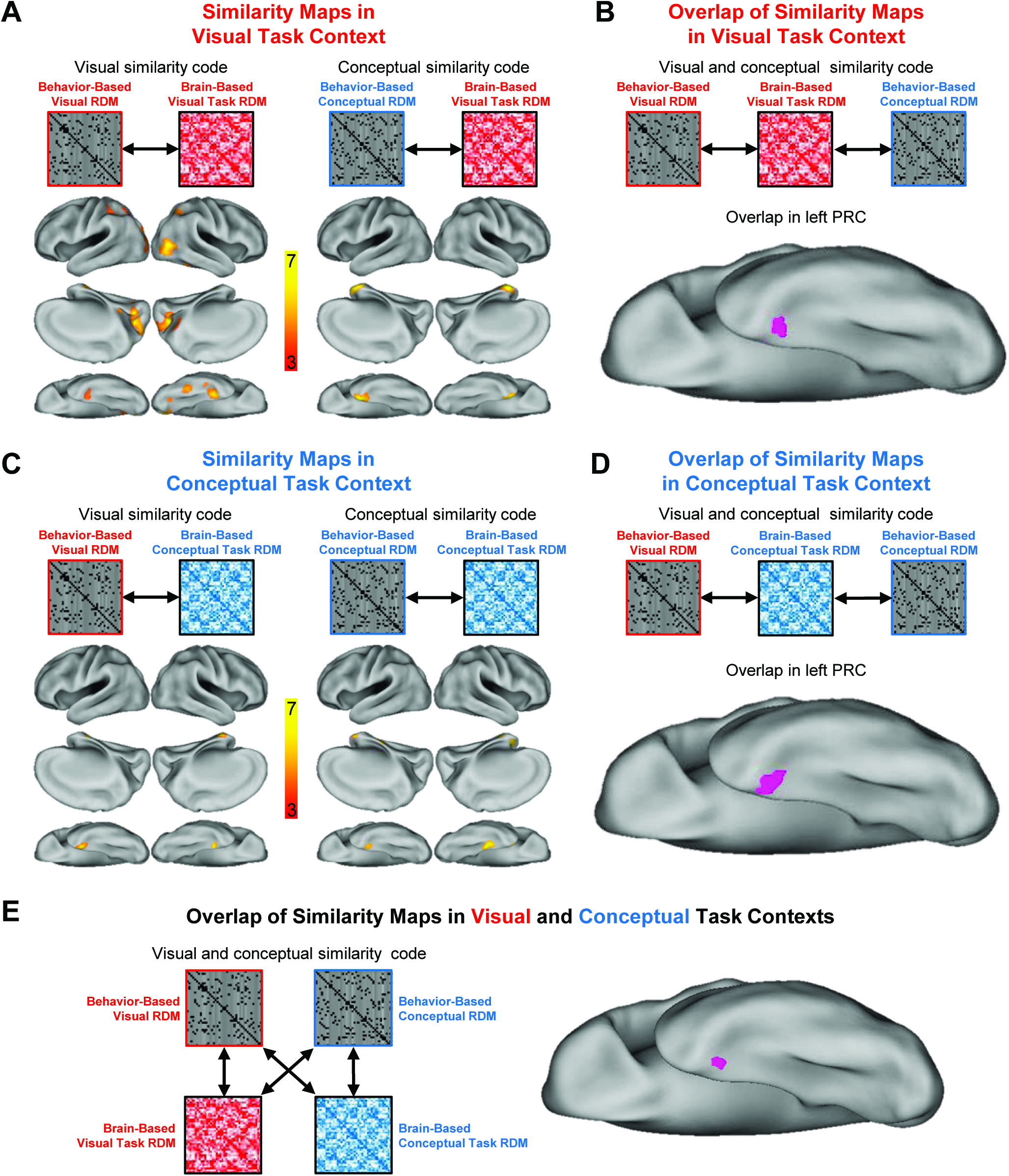
Representational similarity searchlight mapping results. (A) Cortical regions in which the brain-based visual task representational dissimilarity matrix (RDM) was significantly correlated with the behavior-based visual RDM (left) and the behavior-based conceptual RDM (right). **(B)** Overlap between brain-behavior similarity maps in the visual task context. **(C)** Cortical regions in which the brain-based conceptual task RDM was significantly correlated with the behavior-based visual RDM (left) and the behavior-based conceptual RDM (right). **(D)** Overlap between brain-behavior similarity maps in conceptual task context. **(E)** Overlap among brain-behavior similarity maps across both task contexts. The correlation coefficients (Kendall’s tau-a) obtained between behavior-based RDMs and brain-based RDMs were Fisher-*z* transformed and mapped to the voxel at the centre of each searchlight to create the whole-brain similarity maps in panels A and C. Similarity maps in panels A and C were corrected for multiple comparisons using threshold-free cluster enhancement (TFCE) with a corrected statistical threshold of *p* < 0.05 on the cluster level (Smith and Nichols, 2009).

**Table 1.** Clusters in which behavior-based RDMs were significantly correlated with brain-based RDMs as revealed using representational similarity searchlight analyses, with corresponding cluster extent, peak *z*-values, and MNI co-ordinates^1^.

| Region | Cluster Extent | Peak $z$-value | x | y | z |
| --- | --- | --- | --- | --- | --- |
| **Visual Task Context** |  |  |  |  |  |
| *Behavior-Based Visual RDM – Brain-Based Visual Task RDM* |  |  |  |  |  |
| Mid calcarine | 1660 | 5.79 | -2 | -74 | 12 |
| R lateral occipital cortex | 455 | 3.89 | 50 | -66 | 4 |
| R perirhinal cortex | 112 | 3.64 | 34 | -12 | -34 |
| L superior parietal lobule | 110 | 3.21 | -32 | -40 | 44 |
| L perirhinal cortex | 76 | 2.85 | -30 | -12 | -36 |
| R superior parietal lobule | 48 | 2.64 | 38 | -54 | 54 |
| R fusiform gyrus | 45 | 2.77 | 40 | -46 | -20 |
| R precuneus | 29 | 2.66 | 12 | -76 | 48 |
| R Inferior Temporal Gyrus | 9 | 2.52 | 44 | -22 | -28 |
| *Behavior-Based Conceptual RDM – Brain-Based Visual Task RDM* |  |  |  |  |  |
| L Perirhinal Cortex | 368 | 3.96 | -24 | 2 | -38 |
| R Perirhinal Cortex | 232 | 3.26 | 22 | 2 | -36 |
| *Overlap* |  |  |  |  |  |
| L Perirhinal Cortex | 22 |  | -30 | -8 | -38 |
| **Conceptual Task Context** |  |  |  |  |  |
| *Behavior-Based Conceptual RDM – Brain-Based Conceptual Task RDM* |  |  |  |  |  |
| L Perirhinal Cortex | 79 | 2.88 | -30 | -10 | -34 |
| R Parahippocampal Cortex | 64 | 2.94 | 30 | -24 | -24 |
| L Temporal Pole | 61 | 2.89 | -34 | 4 | -26 |
| R Temporal Pole | 25 | 2.70 | 24 | 12 | -36 |
| *Behavior-Based Visual RDM – Brain-Based Conceptual Task RDM* |  |  |  |  |  |
| L Perirhinal Cortex | 98 | 4.87 | -26 | -4 | -10 |
| R Perirhinal Cortex | 26 | 3.01 | 28 | -12 | -34 |
| *Overlap* |  |  |  |  |  |
| L Perirhinal Cortex | 31 |  | -26 | -8 | -42 |
| **Overlap across all Behavior-Based RDMs and Brain-Based RDMs** |  |  |  |  |  |
| L Perirhinal Cortex | 16 |  | -30 | -8 | -36 |

<sup>1</sup>MNI co-ordinates are reported for the peak voxel in individual clusters and the centre of mass for cluster overlap.

In a second step, we examined whether any voxels were common across the task-specific overlapping clusters. In other words, we asked whether both behavior-based RDMs were able to describe the both brain-based RDMs derived from a common set of voxels (as depicted by the black arrows in Figure 6E). Critically, left PRC was the only region in the entire scanned volume in significant clusters of voxels overlapped across all similarity maps (Figure 6E). This result indicates that a subset of voxels within PRC simultaneously expressed both visual and conceptual similarity structure, suggesting that this structure does indeed support integration of the visual and conceptual features that define an object concept.

## Discussion

Although decades of research have aimed at understanding how object concepts are represented in the brain (Warrington et al., 1975; Hodges et al., 1992; Martin et al., 1995; Murray and Bussey, 1999; Chen et al., 2017), the fundamental question of whether and where their conceptual and perceptual features are integrated remains unanswered. Progress toward this end has been hindered by the fact that such features tend to vary concomitantly across object concepts. Here, we used a data-driven approach to systematically select a set of object concepts in which visual and conceptual features varied independently (e.g., hairdryer – comb, which are conceptually but not visually similar; hairdryer – gun, which are visually but not conceptually similar). By comparing behavior-based models of the visual and conceptual similarity structure of these object concepts with corresponding brain-based similarity structure we revealed novel evidence for an integrative coding process that binds conceptual object features with observable perceptual features in a task-invariant manner. This integrative coding, which we uniquely found in PRC, may guide complex behavior through the representation of objects and object concepts at the highest level of specificity. Moreover, we also revealed a representational distinction between PRC and the temporal pole as they relate to semantic memory. Namely, whereas PRC showed evidence of integrative coding across conceptual and visual features, neural activity patterns in the temporal pole were best understood in relation to a purely conceptual code. Taken together, these findings provide a first step toward filling a theoretically important gap in the cognitive neuroscience of semantic memory and object representation, more broadly.

Our central finding is that patterns of activity within PRC reflected both the visual and conceptual similarities between object concepts. We interpret this result as evidence for integration for reasons directly related to our experimental design. First, the behavior-based visual RDM and behavior-based conceptual RDM (i.e., the models) used in the current study were not correlated with one another, indicating that these models accounted for different sources of variability in the relationships among the object concepts. For example, the behavior-based conceptual RDM captured a relationship between “hairdryer” and “comb”, where none existed in the behavior-based visual RDM. Second, and despite the fact that these behavior-based RDMs were orthogonal to one another, they could each be used to describe the brain-based RDMs derived from both the visual and conceptual task contexts. Critically, across our ROI-based RSA and our searchlight analysis, PRC was the only region in which we obtained this pattern of results. At the level of interpretation, the importance of these points is perhaps best illustrated with an example from our experiment. Specifically, our results indicated that while participants made *conceptual* judgments about objects in the fMRI scanner, such as whether a “hairdryer” is man-made or a “gun” is pleasant, the corresponding degree of neural pattern similarity between “hairdryer” and “gun” could be captured by their *perceptual* similarity, as indexed by behavioral ratings from an independent group of observers. Likewise, when participants made *perceptual* judgments about object concepts in the fMRI task, such as whether a “hairdryer” is angular or a “comb” is elongated, the corresponding degree of neural pattern similarity between “hairdryer” and “comb” could be captured by their *conceptual* similarity, as derived from responses provided by an independent group of participants. In both cases, PRC carried information about semantic features that were neither required to perform the immediate task at hand, nor correlated with the features that did in fact have task-relevant diagnostic value. Moreover, results from our RSA-based searchlight mapping analysis indicated that a contiguous cluster of voxels in left PRC was the only region in the brain that showed this effect. Thus, despite the fact that we disentangled conceptual and perceptual feature overlap across objects and imposed task demands that biased processing toward one class of feature or the other, both types of information were ubiquitous and inseparable in PRC. When considered together, these results suggest that, at the level of PRC, it may not be possible to fully disentangle conceptual and perceptual information.

Convergent evidence from studies of functional and structural connectivity in humans, non-human primates, and rodents indicates that PRC is connected to the temporal pole, parahippocampal cortex, LOC, and nearly all other unimodal and polymodal sensory regions in neocortex (Suzuki and Amaral, 1994; Burwell and Amaral, 1998; Kahn et al., 2008; McLelland et al., 2014; Suzuki and Naya, 2014; Wang et al., 2016; Zhuo et al., 2016). Thus, PRC has the connectivity properties that make it well suited to be a multi-modal convergence zone that integrates object features that are both conceptual and perceptual in nature. Indeed, our results have linked LOC to the representation of visual semantic attributes, the temporal pole and parahippocampal cortex to the representation of conceptual attributes, and PRC to the representation of both types of object features. Notably, however, additional research is necessary to directly characterize the nature and direction of semantic information exchanged among these regions. Our findings are also of relevance to the proposal that PRC represents objects in a manner that reflects the highest degree of feature-based integration (i.e., the representational-hierarchical model) (Murray et al., 2007; Graham et al., 2010; Barense et al., 2010, 2011; see Lehky and Tanaka, 2016, for related discussion). Whereas previous research has primarily described its functional role at the level of either visual properties (Buckley and Gaffan, 2006; Murray et al., 2007; Graham et al., 2010; Barense et al., 2012) or semantic attributes (Noppeny et al., 2007; Bruffaerts et al., 2013; Clarke and Tyler, 2014, 2015), here we show for the first time that PRC integrates both types of features, perhaps at the level of fully-specified object representations.

What is the behavioral relevance of highly-specified object representations in which perceptual and conceptual features are integrated? It has previously been suggested that such representations allow for discrimination among stimuli with extensive feature overlap, such as exemplars from the same category (Murray and Bussey, 1999; Noppeny et al., 2007; Graham et al., 2010; Clarke and Tyler, 2015). In line with this view, individuals with medial ATL lesions that include PRC typically have more pronounced conceptual impairments related to living than non-living things (Warrington and Shallice, 1984; Moss et al., 1997, Bozeat et al., 2003), and more striking perceptual impairments for objects that are visually similar as compared to visually distinct (Barense et al., 2007, 2010; Lee et al., 2006). In neurologically healthy individuals, fMRI studies have also demonstrated increased PRC engagement for living as compared to non-living objects (Moss et al., 2005), for known as compared to novel faces (Barense et al., 2011; Peterson et al., 2012), and for faces or conceptually meaningless stimuli with high feature overlap as compared to low (O’Neil et al., 2009; Barense et al., 2012). In a related manner, highly-specified object representations in PRC have also been linked to long-term memory judgments. For example, PRC has been linked to explicit recognition memory judgments when previously studied and novel items are from the same stimulus category (e.g., faces) (Martin et al., 2013, 2016), and when subjects make judgments about their lifetime of experience with a given object concept (Duke et al., 2016). Common among these task demands and experimental manipulations is the requirement to discriminate among highly similar stimuli. In such scenarios, a highly-specified representation that reflects the integration of perceptual and conceptual features necessarily enables more fine-grained distinctions than a purely perceptual or conceptual representation.

This study also has significant implications for prominent neurocognitive models of semantic memory that have characterized the ATL as a semantic hub (Rogers et al., 2006; Patterson et al., 2007; Tranel, 2009). On this view, the bilateral ATLs are thought to constitute a trans-modal convergence zone that abstracts conceptual information from the co-occurrence of features otherwise represented in a distributed manner across modality-specific cortical nodes. Consistent with this idea, we have shown that a behavior-based conceptual similarity model predicted the similarity structure of neural activity patterns in the temporal pole, irrespective of task context. Specifically, neural activity patterns associated with conceptually similar object concepts that are visually distinct (e.g., “hairdryer” – “comb”) were more comparable than were conceptually dissimilar concepts that are visually similar (e.g., “hairdryer” – “gun”), even when task demands required a critical assessment of visual features. This observation, together with results obtained in PRC, demonstrates a representational distinction between distinct ATL structures, a conclusion that dovetails with recent evidence indicating that this region is not functionally homogeneous (Binney et al., 2010; Murphy et al., 2017). Rather, this outcome suggests that some ATL sub-regions play a prominent role in task-invariant extraction of conceptual object properties (e.g., temporal pole), whereas others appear to make differential contributions to the task-invariant integration of perceptual and conceptual features (e.g., PRC) (Lambon Ralph et al., 2017; Chen et al., 2017).

In summary, we used fMRI to characterize the representational structure of object concepts in the brain. Specifically, we generated behavior-based models that independently captured the conceptual and visual similarities among a targeted set of object concepts and used these models to predict brain-based neural similarities across two task contexts. Using this approach we revealed three distinct types of coding of object concepts. First, we found that LOC represented object concepts in a visually-based similarity code. Second, we found that the temporal pole and parahippocampal cortex represented object concepts in a conceptually-based similarity code, but that the temporal pole did so in a task invariant manner, whereas parahippocampal cortex only did so in the context of explicit conceptual feature judgments. Critically, and despite the fact that our visual and conceptual similarity models were not correlated with one another, we found that PRC uniquely supported the integrative coding of perceptual and conceptual features in a task invariant manner. At a broad level, our results suggest that PRC supports the representation of fully-specified object concepts in which perceptual and conceptual information is integrated.

## Methods

### Participants

#### Behavior-Based Visual Similarity Rating Task and Conceptual Feature Generation Task

A total of 2846 individuals completed online behavioral tasks using Amazon’s Mechanical Turk (https://www.mturk.com). Data from 61 participants were discarded due to technical errors, incomplete submissions, or missed catch trials. Of the remaining 2785 participants, 1185 completed the visual similarity rating task (616 males, 569 females; age range = 18-53; mean age = 30.1), and 1600 completed the semantic feature generation task (852 males, 748 females; age range = 18-58 years; mean age = 31.7). Individuals who completed the visual similarity rating task were excluded from completing the feature generation task, and vice versa. All participants provided informed consent and were compensated for their time. Both online tasks were approved by the University of Toronto Ethics Review Board.

#### Brain-Based fMRI Task

A separate group consisting of sixteen right-handed participants took part in the fMRI experiment (10 female; age range = 19-29 years; mean age = 23.1 years). This sample size is in line with extant fMRI studies that have used comparable analytical procedures to test hypotheses pertaining to object representation in the ventral visual stream and ATL (Bruffaerts et al., 2013; Devereaux et al., 2013; Martin et al., 2013, 2016; Clarke and Tyler, 2014; Erez et al., 2016). Due to technical problems, we were unable to obtain data from one experimental run in two different participants. No participants were removed due to excessive motion using a criterion of 1.5mm of translational displacement. All participants gave informed consent, reported that they were native English speakers, free of neurological and psychiatric disorders, and had normal or corrected to normal vision. Participants were compensated $50. This study was approved by the Baycrest Hospital Research Ethics Board.

### Stimuli

As a starting point, we chained together a list of 80 object concepts in such a way that adjacent items in the list alternated between being conceptually similar but visually distinct and visually similar but conceptually distinct (e.g., bullet – gun – hairdryer – comb; bullet and gun are conceptually but not visually similar, whereas gun and hairdryer are visually but not conceptually similar, and hairdryer and comb are conceptually but not visually similar, etc.). Our initial stimulus set was established using the authors’ subjective impressions. The visual and conceptual similarities between all pairs of object concepts were then quantified by human observers in the context of a visual similarity rating task and a conceptual feature generation task, respectively. Results from these behavioral tasks were then used to select 40 object concepts used throughout the current study.

Participants who completed the visual similarity rating task were presented with 40 pairs of words and asked to rate visual similarity between the object concepts to which they referred (Figure 1A). Responses were made using a 5-point scale (very dissimilar, somewhat dissimilar, neutral, somewhat similar, very similar). Each participant was also presented with four catch trials on which an object concept was paired with itself. Across participants, 95.7% of catch trials were rated as being very similar. Data were excluded from 28 participants who did not rate all four catch trials as being at least ‘somewhat similar’. Every pair of object concepts from the initial set of 80 object concepts (3160) was rated by 15 different participants.

We next quantified conceptual similarities between object concepts based on responses obtained in a conceptual feature generation task (Figure 1B), following task instructions previously described by McRae et al. (2005). Each participant was presented with one object concept and asked to produce a list of up to 15 different types of descriptive features, including functional properties (e.g., what it is used for, where it is used, and when it is used), physical properties (e.g., how it looks, sounds, smells, feels, and tastes), and other facts about it, such as the category to which it belongs or other encyclopedic facts (e.g., where it is from). One example object and its corresponding features from a normative database were presented as an example (McRae et al., 2005). Interpretation and organization of written responses were guided by criteria described by McRae et al. (2005). Features were obtained from 20 different participants for each object concept. Data were excluded from 33 participants who failed to list any features. A total of 4851 unique features were produced across all 80 object concepts and participants. Features listed by fewer than 4 out of 20 participants were considered to be unreliable and discarded for the purpose of all subsequent analyses, leaving 723 unique features. This exclusion criterion is proportionally comparable to that used by McRae et al. (2005). On average, each of the 80 object concepts was associated with 10.6 features.

We used a data-driven approach to select a subset of 40 object concepts from the initial 80-item set. These 40 object concepts are reflected in the behavior-based visual and conceptual RDMs, and were used as stimuli in our fMRI experiment. Specifically, we first ensured that each object concept was visually similar, but conceptually dissimilar, to at least one other item (e.g., hairdryer – gun), and conceptually similar, but visually dissimilar, to at least one different item (e.g., hairdryer – comb). Second, in an effort to ensure that visual and conceptual features varied independently across object concepts, stimuli were selected such that the corresponding behavior-based visual and conceptual similarity models were not correlated with one another.

### Behavior-Based RDMs

#### Behavior-Based Visual RDM

A behavior-based model that captured visual dissimilarities between all pairs of object concepts included in the fMRI experiment (40 object concepts) was derived from the visual similarity judgments obtained from our online rating task. Specifically, similarity ratings for each pair of object concepts were averaged across participants, normalized, and expressed within a 40x40 RDM (1 – averaged normalized rating). Thus, the value in a given cell of this RDM reflects the visual similarity of the object concepts at that intersection. This behavior-based visual RDM is our visual dissimilarity model.

#### Behavior-Based Conceptual RDM

A behavior-based model that captured conceptual dissimilarities between all pairs of object concepts included in the fMRI experiment was derived from data obtained in our online feature-generation task. In order to ensure that the semantic relationships captured by our conceptual similarity model were not influenced by verbal descriptions of visual attributes, we systematically removed features that characterized either visual form or color (e.g., “is round” or “is red”). Using these criteria a total of 58 features (8% of the total number of features provided) were removed. We next quantified conceptual similarity using a concept-feature matrix in which rows corresponded to object concepts (i.e., 40 rows) and columns to conceptual features (i.e., 723 features – 58 visual features = 665 columns) (Figure 1B, center). Specifically, we computed the cosine angle between each row; cosine similarity reflects the conceptual distances between object concepts such that high cosine similarities between items denote short conceptual distance. The conceptual dissimilarities between all pairs of object concepts were expressed as a 40 x 40 RDM. The value within each cell of the conceptual model RDM was calculated as 1 – the cosine similarity value between the corresponding object concepts. This behavior-based conceptual RDM is our conceptual dissimilarity model.

#### Behavior-Based RSA: Comparison of Behavior-Based RDMs

We next quantified similarity between our behavior-based visual RDM and behavior-based conceptual RDM using Kendall’s tau-a as the relatedness measure. This ranked correlation coefficient is the most appropriate inferential statistic to use when comparing sparse RDMs that predict many tied ranks (i.e., both models predict complete dissimilarity between many object pairs; Nili et al., 2014). Inferential analysis of model similarity was performed using a stimulus-label randomization test (10,000 iterations) that simulated the null hypothesis of unrelated RDMs (i.e., zero correlation) based on the obtained variance. Significance was assessed through comparison of the obtained Kendall’s tau-a coefficient to the equivalent distribution of ranked null values. As noted in the Results section, this analysis revealed that our behavior-based visual and conceptual RDMs were not significantly correlated (Kendall’s tau-a = .01, *p* = .09). Moreover, inclusion of the 58 features that described color and visual form in the behavior-based conceptual RDM did not significantly alter its relationship with the visual behavior-based visual RDM (Kendall’s tau-a = .01, *p* = .09).

### Experimental Procedures: fMRI Feature Verification Task

During scanning, participants completed a feature verification task that required a yes/no judgment indicating whether a given feature was applicable to a specific object concept on a trial-by-trial basis. We systematically varied the feature verification probes in a manner that established a visual feature verification task context and conceptual feature verification task context. Verification probes comprising the visual task context were selected to encourage processing of the visual semantic features that characterize each object concept (i.e., shape, color, and surface detail). To this end, eight specific probes were used: shape [(angular, rounded), (elongated, symmetrical)], color (light, dark), and surface (smooth, rough). Notably, all features are associated with two opposing probes (e.g., angular and rounded; natural and manufactured) to ensure that participants made an equal number of “yes” and “no” responses. Verification probes comprising the conceptual feature verification task context were selected to encourage processing of the abstract conceptual features that characterize each object concept (i.e., animacy, origin, function, and affective associations). To this end, eight specific verification probes were used: (living, non-living), (manufactured, natural), (tool, non-tool), (pleasant, unpleasant).

#### Procedures

The primary experimental task was evenly divided over eight runs of functional data acquisition. Each run lasted 7m 56s and was evenly divided into two blocks, each of which corresponded to either a visual verification task context or a conceptual feature verification task context. The order of task blocks was counter-balanced across participants. Each block was associated with a different feature verification probe, with the first and second block in each run separated by 12s of rest. Blocks began with an 8s presentation of a feature verification probe that was to be referenced for all intra-block trials. With this design, each object concept was repeated 16 times: eight repetitions across the visual feature verification task context and eight repetitions across the conceptual feature verification task context. Behavioral responses were recorded using an MR-compatible keypad.

Stimuli were centrally presented for 2s and each trial was separated by a jittered period of baseline fixation that ranged 2-6s. Trial order and jitter interval were optimized for each run using the OptSeq2 algorithm(http://surfer.nmr.mgh.harvard.edu/optseq/), with unique sequences and timing across counterbalanced versions of the experiment. Stimulus presentation and timing was controlled by E-Prime 2.0 (Psychology Software Tools, Pittsburgh, PA).

### Experimental Procedure: fMRI Functional Localizer Task

Following completion of the main experimental task, each participant completed an independent functional localizer scan that was subsequently used to identify LOC. Participants viewed objects, scrambled objects, words, scrambled words, faces, and scenes in separate 24s blocks (12 functional volumes). Within each block, 32 images were presented for 400ms each with a 350ms ISI. There were four groups of six blocks, with each group separated by a 12s fixation period, and each block corresponding to a different stimulus category. Block order (i.e., stimulus category) was counterbalanced across groups. All stimuli were presented in the context of a 1-back task to ensure that participants remained engaged throughout the entire scan. Presentation of images within blocks was pseudo-random with 1-back repetition occurring 1-2 times per block.

### ROI Definitions

We performed RSA in four a priori defined ROIs. The temporal pole, PRC, and parahippocampal cortex were manually defined in both the left and right hemisphere on each participant’s high-resolution anatomical image according to established MR-based protocols (Pruessner et al., 2002, with adjustment of posterior border of parahippocampal cortex using anatomical landmarks described by Frankó et al., 2014). Lateral occipital cortex was defined as the set of contiguous voxels located along the lateral extent of the occipital lobe that responded more strongly to intact than scrambled objects (*p* < 0.001, uncorrected; Malach et al. 1995).

### fMRI Data Acquisition

Scanning was performed using a 3.0-T Siemens MAGNETOM Trio MRI scanner at the Rotman Research Institute at Baycrest Hospital using a 32-channel receiver head coil. Each scanning session began with the acquisition of a whole-brain high-resolution magnetization-prepared rapid gradient-echo T1-weighted structural image (repetition time = 2s, echo time = 2.63ms, flip angle = 9°, field of view = 25.6cm^2^, 160 oblique axial slices, 192 × 256 matrix, slice thickness = 1mm). During each of eight functional scanning runs comprising the main experimental task, a total of 238 T2*-weighted echo-planar images were acquired using a two-shot gradient echo sequence (200 × 200 mm field of view with a 64 × 64 matrix size), resulting in an in-plane resolution of 3.1 × 3.1 mm for each of 40 2-mm axial slices that were acquired in an interleaved manner along the axis of the hippocampus. The inter-slice gap was 0.5 mm; repetition time = 2s; echo time = 30ms; flip angle = 78°). These parameters yielded coverage of the majority of cortex, excluding only the most superior aspects of the frontal and parietal lobes. During a single functional localizer scan, a total of 360 T2*-weighted echo-planar images were acquired using the same parameters reported for the main experimental task. Lastly, a B0 field map was collected following completion of the functional localizer scan

### fMRI Data Analysis Software

Preprocessing and GLM analyses were performed in FSL5 (Smith et al., 2004). Representational similarity analyses were performed using CoSMoMVPA (http://www.cosmomvpa.org/; Oosterhof et al., 2016) together with custom Matlab code (The MathWorks, Inc., Natick, MA).

### Preprocessing and Estimation of Object-Specific Multi-Voxel Activity Patterns

Images were initially skull-stripped using a brain extraction tool (BET, Smith, 2002) to remove non-brain tissue from the image. Data were then corrected for slice-acquisition time, high-pass temporally filtered (using a 50s period cut-off for event-related runs, and a 128s period cut-off for the blocked localizer run), and motion corrected (MCFLIRT, Jenkinson et al., 2002). Functional runs were registered to each participant’s high-resolution MPRAGE image using FLIRT boundary-based registration with B0-fieldmap correction. The resulting unsmoothed data were analyzed using first-level FEAT (v6.00; fsl.fmrib.ox.ac.uk/fsl/fslwiki) in each participant’s native anatomical space. Parameter estimates of BOLD response amplitude were computed using FILM, with a general linear model that included temporal autocorrelation correction and 6 motion parameters as nuisance covariates. Each trial (i.e., object concept) was modeled with a delta function corresponding to the stimulus presentation onset and then convolved with a double-gamma hemodynamic response function. Separate response-amplitude (β) images were created for each object concept (n = 40), in each run (n = 8), in each property verification task context (n = 2). Obtained β images were converted into *t*-statistic maps; previous research has demonstrated a modest advantage for *t*-maps over β images in the context of multi-voxel pattern analysis (Misaki et al., 2010). These data were used for all subsequent similarity analyses.

### Representational Similarity Analysis (RSA)

#### ROI-Based RSA: Comparisons of Behavior-Based RDMs with Brain-Based RDMs and Brain-Based RDMs with Brain-Based RDMs

We used linear correlations to quantify the participant-specific dissimilarities (1 - Pearson’s r) between all object-evoked multi-voxel activity patterns (n = 40) with each ROI (n = 4). Dissimilarity measures were expressed in 40x40 RDMs for each run (n = 8) and verification task context (n = 2), separately. Thus, for each ROI, each participant had eight RDMs that reflected the (dis)similarity structure from the visual feature verification task context, and eight RDMs that reflected the (dis)similarity structure from the conceptual verification task context. We then calculated one mean RDM for each feature verification task context by averaging run-specific RDMs across participants. Thus, one brain-based RDM was created for the visual task context (i.e., brain-based visual task RDM) and one brain-based RDM was created for the conceptual task context (i.e., brain-based conceptual task RDM).

We next examined how well each of the behavior-based RDMs fit each of the obtained brain-based RDMs for each ROI. Model fit was quantified as the ranked correlation coefficient (Kendall’s tau-a) between behavior-based RDMs and the brain-based RDMs. Significance testing was performed using a stimulus-label randomization test (10,000 iterations per model) Bonferroni corrected for multiple comparisons.

#### Searchlight-Based RSA

Whole-volume RSA was implemented using 100-voxel surface-based searchlights (Kriegeskorte et al., 2006; Oosterhof et al., 2011). Each surface-based searchlight referenced the 100 nearest voxels to the searchlight center based on geodesic distance on the cortical surface. Neural estimates of dissimilarity (i.e., RDMs) were calculated in each searchlight using the same approach implemented in our ROI-based RSA. Correlations between behavior-based RDMs were also quantified using the same approach. The correlation coefficients obtained between behavior-based RDMs and brain-based RDMs were then Fisher-*z* transformed and mapped to the voxel at the centre of each searchlight to create a whole-brain similarity map. Participant-specific similarity maps were then normalized to a standard MNI template using FNIRT (Greve and Fischl, 2009). To assess the statistical significance of searchlight maps across participants, all maps were corrected for multiple comparisons without choosing an arbitrary uncorrected threshold using threshold-free cluster enhancement (TFCE) with a corrected statistical threshold of *p* < 0.05 on the cluster level (Smith and Nichols, 2009). A Monte Carlo simulation permuting condition labels was used to estimate a null TFCE distribution. First, 100 null searchlight maps were generated for each participant by randomly permuting condition labels within each obtained searchlight RDM. Next, 10,000 null TFCE maps were constructed by randomly sampling from these null data sets in order to estimate a null TFCE distribution (Stelzer et al., 2013). The resulting surface-based statistically thresholded *z*-score were projected onto the PALS-B12 surface atlas in CARET version 5.6. (http://www.nitrc.org/projects/caret/; Van Essen et al., 2001; Van Essen, 2005).

## Acknowledgements

This work was supported by the Canadian Natural Sciences Engineering Research Council (Discovery and Accelerator Grants to M.D.B.; Postdoctoral Fellowship award to C.B.M.), the James S. McDonnell Foundation (Scholar Award to M.D.B.), and the Canada Research Chairs Program (M.D.B.).

## Footnotes

1 MNI co-ordinates are reported for the peak voxel in individual clusters and the centre of mass for cluster overlap.

